# Automated Detection and Diameter Estimation for Mouse Mesenteric Artery using Semantic Segmentation

**DOI:** 10.1101/2020.10.28.358721

**Authors:** Akinori Higaki, Ahmad U. M. Mahmoud, Pierre Paradis, Ernesto L. Schiffrin

## Abstract

**Background:** Pressurized myography is useful for the assessment of small artery structure and function, and widely used in the field of cardiovascular research. However, this procedure requires technical expertise for the sample preparation and effort to choose an appropriate size of artery. In this study we sought to develop an automatic artery-vein differentiation and size measurement system utilizing the U-Net-based machine learning algorithms.

**Methods and Results:** We used 654 independent mesenteric artery images from 59 mice for the model training and validation. Our segmentation model yielded 0.744 ±0.031 in IoU and 0.881 ±0.016 in Dice coefficient with 5-fold cross validation. The vessel size and the lumen size calculated from the predicted vessel contours demonstrated a strong linear correlation with the manually determined vessel sizes (R = 0.722 ±0.048, p<0.001 for vessel size and R = 0.908 ±0.027, p<0.001 for lumen size). Lastly, we assessed the relation between the vessel size before and after dissection using pressurized myography system. We observed a strong positive correlation between the wall/lumen ratio before dissection and the lumen expansion ratio (R^2^ = 0.671, p<0.01). Using multivariate binary logistic regression, two models estimating whether the vessel met the size criteria (lumen size of 160 to 240 μm) were generated with area under the ROC curve of 0.761 for the upper limit and 0.747 for the lower limit.

**Conclusion:** Our novel image analysis method with U-Net could streamline the experimental approach and may facilitate cardiovascular research.

## 1. INTRODUCTION

Since the dysfunction of resistance vessels contributes to blood pressure elevation (1,2) and is an independent risk factor for cardiovascular events (3), a number of previous reports have focused on the study of small mesenteric artery structure and function in hypertension and cardiovascular disease (4,5). Pressurized myography is one of the most useful tools for the assessment of small arteries and widely used in cardiovascular research (6,7). However, this procedure requires technical expertise for the sample preparation and takes effort and training to choose an appropriate size artery (8). Therefore, automatic vessel detection with a size measurement system could be helpful to avoid unnecessary work on inappropriate vessels, increasing the success rate of experiments.

Along with recent advances in computational science, machine-learning methods including deep learning have been increasingly used in medical research (9). Recently, Hemelings et al. reported a method for artery/vein differentiation in retinal images using a fully convolutional neural network (FCN) with U-Net architecture (10). Another study showed the utility of U-Net for the size prediction of cerebral aneurysms in magnetic resonance angiography (11). Based on our experience with pressurized myography, we hypothesized that the appropriateness of an artery for experimentation with respect to its diameter can be estimated based on the parameters before dissection, which are calculated from the contours obtained by the segmentation model.

Here, we propose a novel workflow for image segmentation of mouse mesenteric arteries using U-Net, and demonstrate the validity of our size measurement algorithm for vessel diameters before dissection and prediction of vessel size after pressurization at 45 mm Hg.

## 2. MATERIAL AND METHODS

Datasets including script files to reproduce these results are available from the online repository (http://dx.doi.org/10.17632/6swz3x26y3.1).

### 2.1 Image acquisition from animal samples

The study was approved by the Animal Care Committee of the Lady Davis Institute for Medical Research and McGill University and followed recommendations of the Canadian Council of Animal Care. Six hundred and fifty-four 2^nd^ and 3^rd^ order mesenteric artery images from 59 mice, which were already destined to be euthanized for other research purposes, were collected from September 2019 to April 2020 (Supplementary Table). This included wild-type C57BL/6J, wild-type C57BL/6NHsd, ieCre, ieET-1, eET-1, eET-1, *ApoE*^-/-^, *eET-1*/*Apoe*^-/-^, *Apoe*^+/-^/*Nox1*^-/y^, *Apoe*^-/-^/*Nox1*^-/y^, *Apoe*^-/-^/*Nox4*^+/-^, eET-1/*Apoe*^+/-^/*Nox1*^-/y^, eET-1/*Apoe*^-/-^/*Nox1*^-/y^, eET-1/*Apoe*^-/-^/*Nox4*^+/-^, *Il-23r^GFP/GFP^*, and *P2rx7*^-/-^ mice. All the mice, except one wildtype C57BL/6NHsd mouse and 3 ieCre mice, were male. The wild-type C57BL/6J, *Apoe*^-/-^ (apolipoprotein E knockout) and *P2rx7*^-/-^ (P2X purinoceptor 7 knockout) mice were obtained from Jackson Laboratories (Bar Harbor, ME, USA) and the wild-type C57BL/6NHsd mice from Envigo (Indianapolis, IN, USA). The eET-1 mice were generated in our laboratory (12) and are C57BL/6NHsd transgenic mice overexpressing the human endothelin (ET)-1 driven by the *Tie2* promoter conferring endothelial-specific expression. *Nox1*^-/y^ (*Nox1* knockout) and *Nox4*^-/-^ (*Nox4* knockout) mice are NADPH oxidase 1 and 4 knockout have been described previously (13,14). The *Apoe*^-/-^/*Nox1*^-/y^ and *Apoe*^-/-^/*Nox4*^-/-^ mice were obtained from Dr. Karin A. Jandeleit-Dahm at the Baker IDI Heart & Diabetes Research Institute (Melbourne, Australia) (15). The eET-1/*Apoe*^-/-^, eET-1/*Apoe*^-/-^/*Nox1*^-/y^ and eET-1/*Apoe*^-/-^/*Nox4*^-/-^ mice were generated at the Lady Davis Institute for Medical Research (Montreal, QC, Canada) by crossing eET-1 mice with *Apoe^-/-^* mice, and then eET-1/*Apoe*^-/-^ with *Apoe*^-/-^/*Nox1*^-/y^ or *Apoe*^-/-^/*Nox4*^-/-^ mice. The ieCre (also known as Tie2-CreER^T2^) mice are transgenic mice expressing a tamoxifen-inducible Cre recombinase (16) under the control of the endothelium-specific *Tie2* promoter (17). The ieET-1 mice were generated in our laboratory (18) and are tamoxifen-inducible endothelium-restricted human ET-1 overexpressing transgenic mice. The male ieCre and ieET-1 mice were treated with tamoxifen (1 mg/day, S.C.) for 5 days to induce the Cre recombinase at 9-12 weeks old and studied 3 months later. The *Il-23r^GFP/GFP^* mice were provided by Dr. Mohamed Oukka (Department of Immunology, University of Washington, Seattle, WA, USA) and are knock-in mice that have the intracellular domain of the interleukin 23 receptor (IL-23R) replaced by the green fluorescent protein (GFP) (19). A subset of wild-type C57BL/6J and *P2rx7^-/-^* mice were infused with angiotensin II subcutaneously (1000 ng/kg/min) for 14 days with ALZET osmotic mini pumps (Model 1002, Durect Corporation, Cupertino, CA) (Supplementary Table).

Image acquisition and the following annotation task was carried out by two independent investigators. On the day of the study, mice were anesthetized with 3% isoflurane mixed with O_2_ at 1 L/min, depth of anesthesia confirmed by rear foot squeezing, the mesenteric artery vascular bed attached to the intestine was dissected from the abdominal cavity and collected on ice-cold phosphate-buffered saline (PBS) or Krebs solution (pH 7.4) containing (mmol/l): 120 NaCl, 25 NaHCO_3_, 4.7 KCl, 1.18 KH_2_PO_4_, 1.18 MgSO_4_, 2.5 CaCl_2_, 0.026 EDTA and 5.5 glucose. The mesenteric artery vascular bed was exposed by spreading delicately the intestine on a layer of black wax using pins in a petri dish containing PBS or Krebs. The upper surface of each mesenteric artery was then exposed by dissecting out perivascular adipose tissue. Images were captured using a surgical microscope Leica M651 (Leica Microsystems Inc., Richmond Hill, ON, Canada) at 40x magnification with an Infinity 1-3C camera (Lumenera Co., Ottawa, Canada) at 640x480 resolution. The images were center-cropped to 480x480 pixel size and saved in RGB color JPG format. The images were calibrated using an American Optical AO Microscope Stage 2 mm micrometer with 0.01 mm division (American Optical Company, Buffalo, NY, USA). A single pixel represents 3.154 μm.

A principal component analysis (PCA) was performed for the characterization of the collected image. Each image data was transformed into a single vector consists of 230,400 (480x480) pixel values before calculating the principal components. We used the first and the second components for the 2-dimensional scatter plot. Scikit-learn, a Python library, was used for this analysis.

### 2.2 Dataset preparation

Segmented mask images were made by using Labelme: A Python-based image annotation tool (https://github.com/wkentaro/labelme). Each pixel was labelled into four classes: 1. background, 2. arterial lumen, 3. arterial wall (includes adventitia, media and intima), and 4. off-target vessel such as vein or capillaries on the surface of the intestine. Arteries and veins were differentiated based on the morphology and haptic feedback as described previously (8). Basically, a vein lacks thick media and has a flatter shape compared to the accompanying artery, leading to more transparent appearance and less elasticity on touch. In case of difficulty discriminating arteries from veins, the vessel was classified as off-target. After annotation, all images were saved as indexed color in PNG format, and 20% were left out for 5-fold cross-validation. Besides the segment information, we manually measured vessel diameters using NIH ImageJ software (https://imagej.nih.gov/ij/index.html) at the center of the artery.

### 2.3 Segmentation model

We employed U-Net architecture as a semantic segmentation model (20) and made some modification fit to our needs. We unified the sizes of input and output image as 480x480 by padding after the convolution operation, while the original implementation uses cropping from different sizes. To shorten the processing time, we used lower number of kernels in each layer compared to the original implementation (24 channels in the first convolution layer instead of 36). We employed the Dice coefficient as a loss function, instead of cross entropy, on the basis of reports that using Dice coefficient is more efficient in the segmentation task when the targeted object is relatively small compared to the background object (21,22). A schematic diagram of the FCN model is shown in **Figure 1**. Segmented images were used as ground truth to train the machine learning model. Simple geometric data augmentation (90-degree clockwise rotation) was applied to the training dataset, based on the assumption that there were two types of investigators: those who were more comfortable working with vessels in vertical orientation and those who preferred a horizontal orientation.

**Figure 1.**
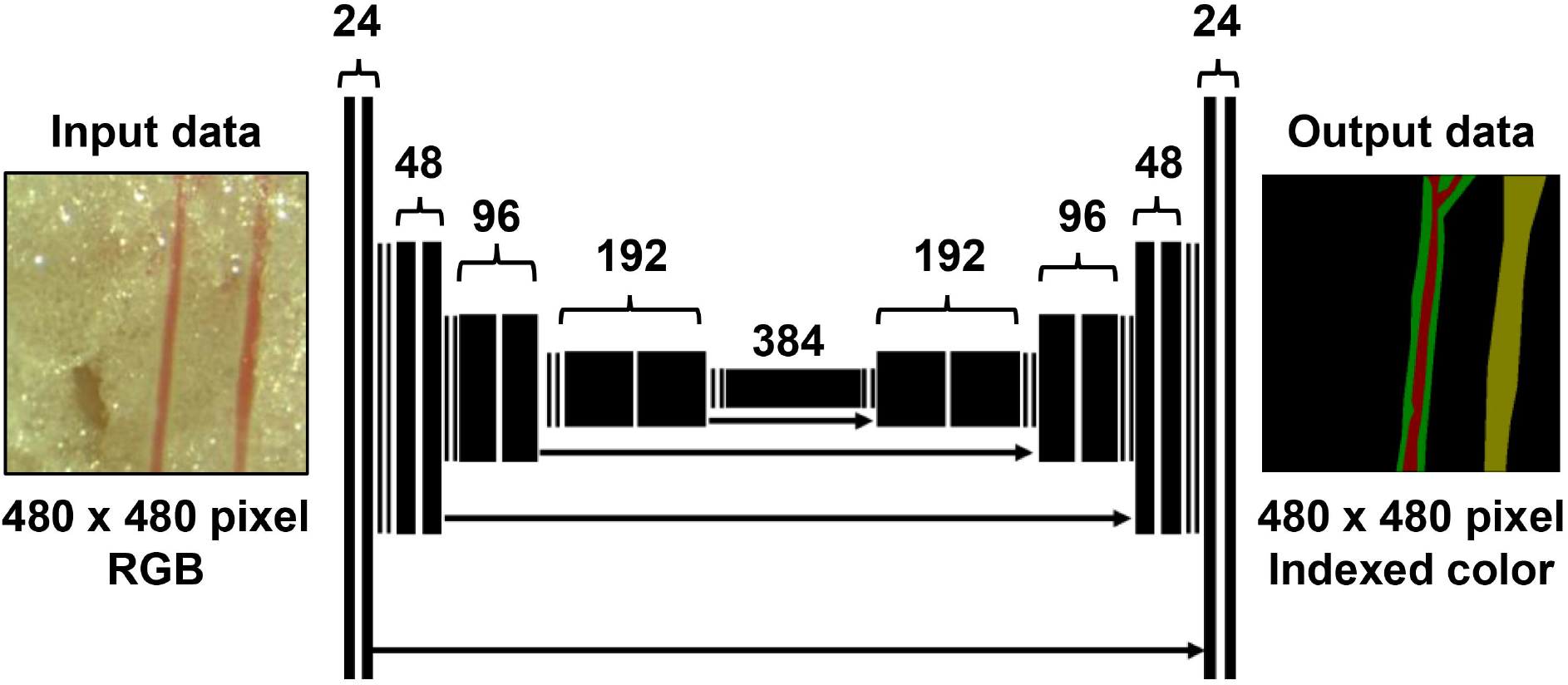
Schematic representation of the U-Net-based segmentation model. Each black box corresponds to a multi-channel feature map. The number of channels is denoted above the boxes. The arrows denote contracting paths.

### 2.4 Algorithm for the automated linewidth measurement

From the segmented output images, the vessel area and lumen area were obtained as binary images. Using OpenCV python library (https://pypi.org/project/opencv-python/), total circumference length of the vessel contour and lumen contour were obtained respectively. Based on rectangular approximation, the circumference (denoted as *C*) was calculated as *C* = 2(*w + h*), where *w* and *h* are average width and height of the assumptive rectangles. Each rectangle area was also approximated as *A* = *w × h*. By solving these equations, the average vessel and lumen diameter (width) was obtained as 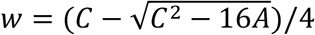 as illustrated in the Supplementary Figure S1. The agreement between the manual and the automated method was evaluated for the outer diameter, lumen diameter and W/L ratio in all image data. This automated measurement system (combined with the automatic segmentation model) was implemented as a publicly available webbased application (https://lda-amma.herokuapp.com).

### 2.5 Evaluation of the model performance on the segmentation task

The model performance for correct segmentation was evaluated based on pixel accuracy, Intersection-over-Union (IoU) and Dice coefficient (alternatively known as F1-score) with 5-fold cross validation. As for the automated diameter measurement, the correlation between the measurement in predicted segmentation and true segmentation was also assessed.

### 2.6 Assessment of vessel diameter in the pressurized myography chamber

Among all 59 animals (654 images) studied, pressurized myography data were obtained from a subset of 35 animals (240 images). The mesenteric artery lumen diameter and the wall thickness of the vessel were measured before dissection as described above. Thereafter, vessels were mounted in the myography chamber and equilibrated for 60 min at 45 mm Hg of intraluminal pressure in PBS or Krebs solution at room temperature. Media and lumen diameters were measured by a video dimension analyzer (Living Systems Instrumentation, Burlington, Virginia, USA) at 45 mm Hg intraluminal pressure. The experimental conditions are determined based on the methodology of our previous reports (7).

### 2.7 Statistical analysis

Data are presented as means ±standard error. Pearson’s correlation was employed for regression analysis, and Pearson correlation coefficient *R*-values higher than 0.7 (therefore 0.49 for *R*^2^) were considered to show strong linear relationships. *P*<0.05 were considered statistically significant. The SciPy module in the Python library was used for these analyses. A multivariate binary logistic regression analysis and receiver operator characteristics (ROC) analysis were performed using IBM SPSS Statistics 24 software for Windows (SPSS, Inc., Chicago, IL, USA) to predict whether the vessel lumen size was within the inclusion criteria: between 160 μm to 240μm at 45mm Hg intraluminal pressure. We made two models that predict 1) whether the vessel size is over 240μm and 2) whether the vessel size in under 160μm. The vessel size, lumen size, wall/lumen ratio determined within the spread mesenteric artery tree and the mouse age were considered as possible explanatory variables. A stepwise method was used to select the explanatory variables for each prediction model based on Wald statistics.

## 3. RESULTS

### 3.1 Characteristics of collected data (descriptive analysis)

The mean age of the mice at the time of study was 14.9 ±0.7 weeks, and 11.4 ±0.9 images were obtained from each mouse. Forty-four % of the mice were wild-type and the rest were genetically engineered (**Supplementary Table**). The majority of the arteries (81%) were captured in vertical orientation and the others were horizontal or multi-oriented. Fifty-nine % of the arteries were captured with the accompanying vein. Seventeen % of the arteries had a bifurcation or side branches. Only 3.2% of the images contain more than two arteries. Fifty-four % of the arteries were surrounded by perivascular adipose tissue. Twelve % of the lumens had lost the blood from their lumen and appeared transparent. Ten % of the arteries were accidentally damaged while handling and had some local dilatation of the lumen or disruption of the wall with bleeding. Examples of collected images are shown with principal components in **Supplementary Figure S2**.

### 3.2 Model performance for segmentation

The average pixel accuracy was 0.942 ±0.007. The average IoU and Dice coefficient were 0.744 ±0.031 and 0.881 ±0.016 respectively. Representative input and output images are shown in **Figure 2**.

**Figure 2.**
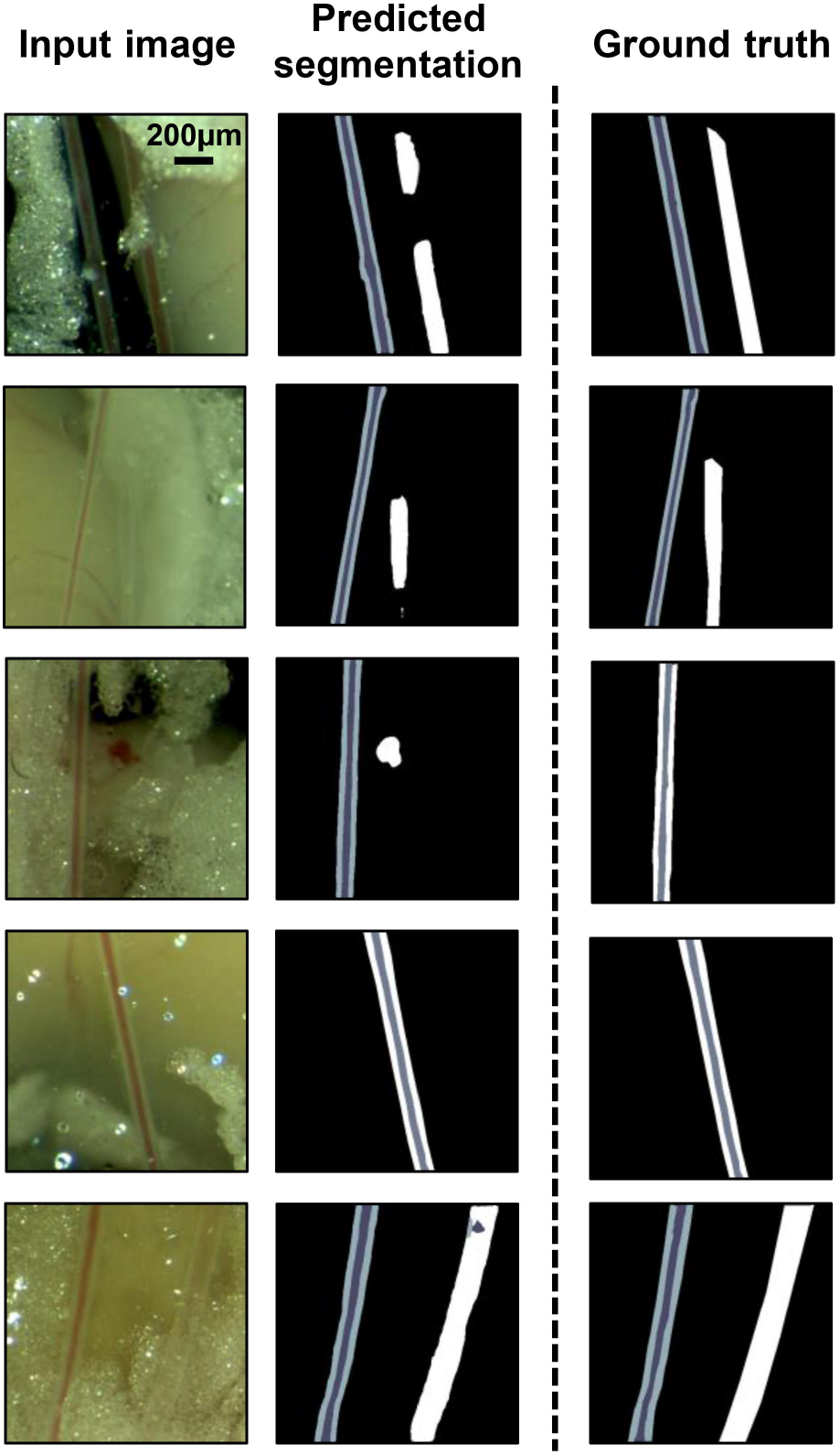
Representative input and output images of the segmentation model. The black, dark gray, light gray and white colors correspond to background, arterial wall, arterial lumen and vein respectively in the segmented images.

### 3.3 Validation of the automated method of line-width calculation

Before applying to the predicted vessel contours, we tested the validity of our rectangular-approximation method for linewidth calculation, by comparing it to the manual measurement. For both vessel size (outside diameter) and lumen size (inner diameter), the automated calculation gave similar values to the manual measurement (R^2^ = 0.831 for vessel size and R^2^ = 0.859 for lumen diameter, *P*<0.001 for both) as shown in **Supplementary Figure S3**.

### 3.4 Automated diameter measurement based on the predicted vessel contour

For both vessel size and lumen size, there was a strong significant linear correlation between the true dimension and the estimated size based on the predicted contour (Supplementary Figure S4). The R value was 0.722 ±0.048 for vessel size and 0.908 ±0.027 for lumen size (*P*<0.001 throughout all the trials of 5-fold cross validation). The representative size estimation outcome is shown in **Figure 3**.

**Figure 3.**
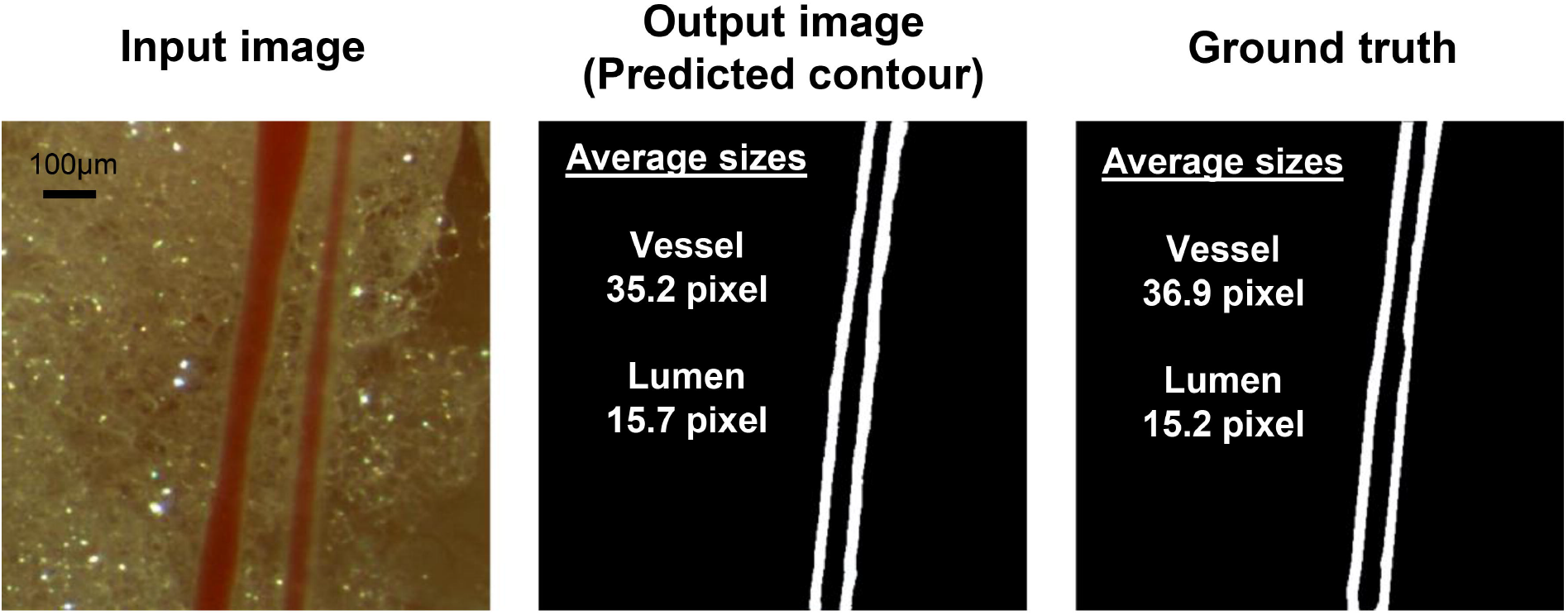
An example of automated vessel size estimation. The vein located on the right side of the artery (Input image) was successfully ignored in the segmented image (Predicted contour) and the contour looked identical to the true contour manual segmentation (Ground truth). The chart shows the line widths obtained by the rectangular-approximation method.

### 3.5 Artery dimensional changes after pressurization at 45 mmHg

Vessel contour before dissection and after pressurization at 45 mm Hg were assessed using 221 vessels from 29 animals. An example image of an artery before and after dissection is shown in **Figure 4**. There was no strong linear correlation of outer vessel or lumen size between before and after pressurization (**Figure 5 A and B**). W/L ratio before dissection had no linear correlation with that at 45mmHg (**Figure 5C**). On the other hand, there was a strong and significant correlation between W/L ratio before dissection and the fold-change of lumen diameter (R^2^ = 0.671, *P*<0.001) as shown in **Figure 5D**.

**Figure 4.**
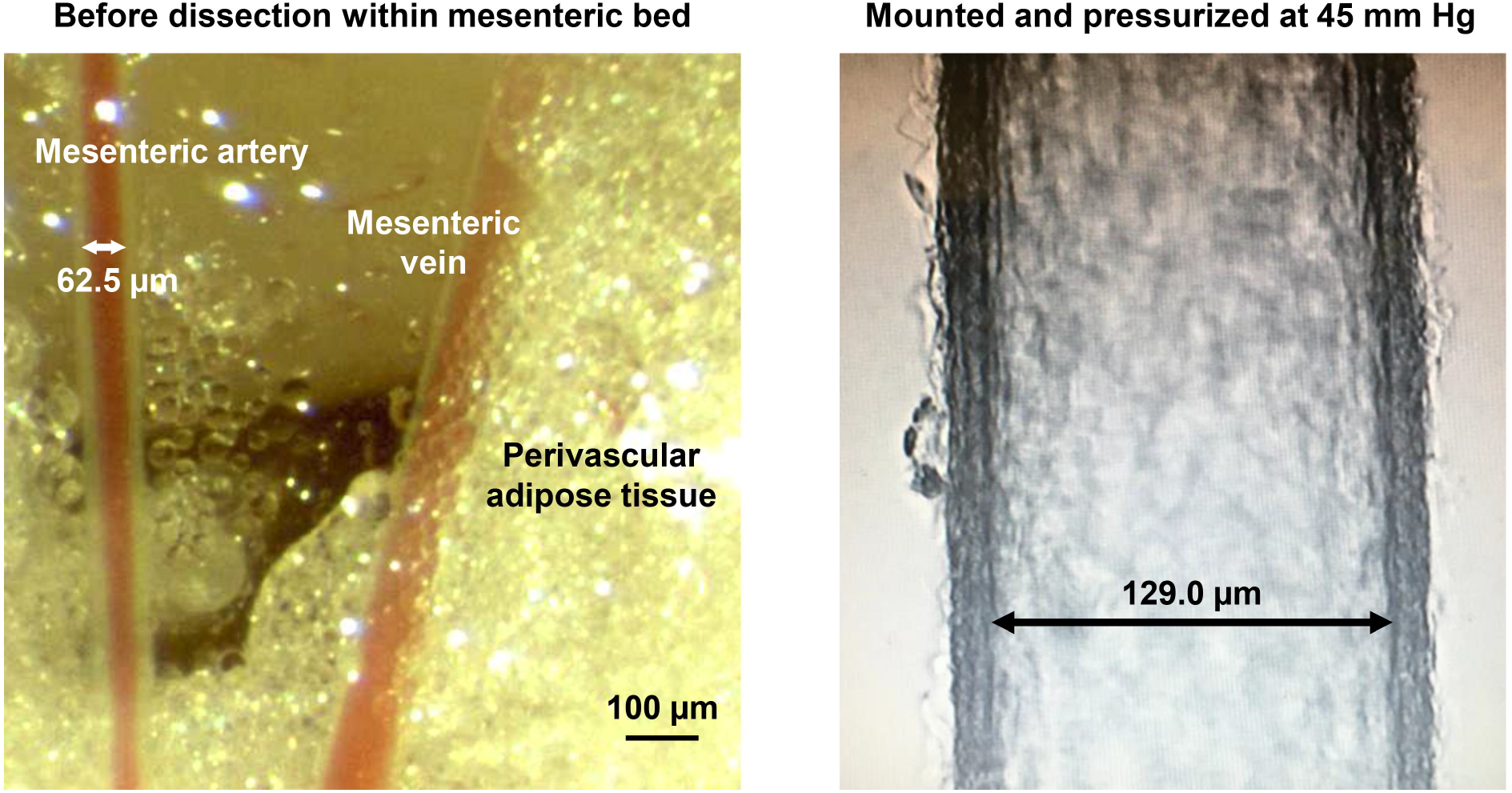
An example image of the artery before and after pressurization at 45 mm Hg. Lumen diameter is indicated in each image. In this case, the fold change of the lumen diameter is 2.06 (=129.0 μm/62.5 μm).

**Figure 5.**
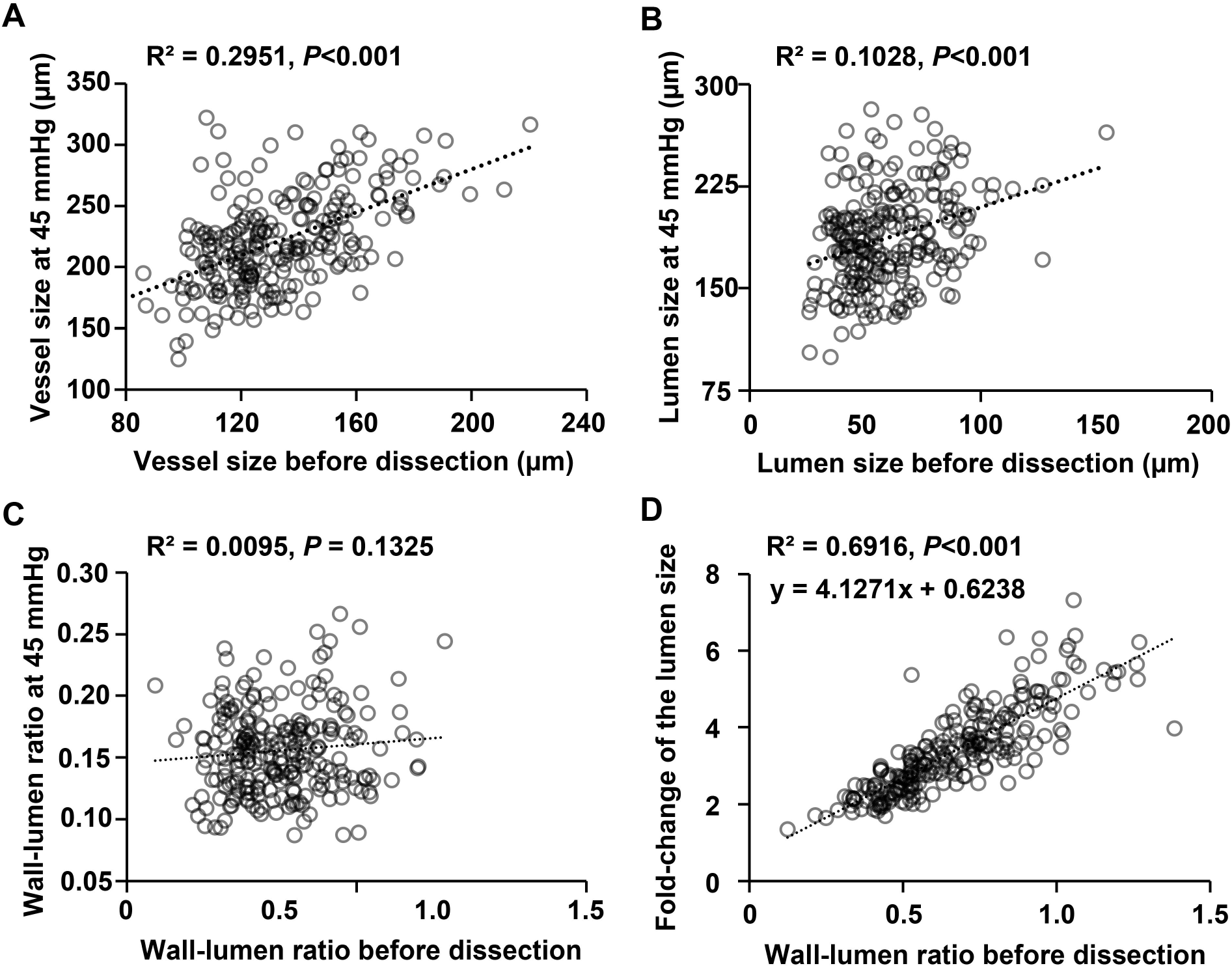
Correlation of dimensional parameters before dissection and after being pressurized at 45mmHg. Scatter plots showing the relation between before dissection and after pressurization are shown for the vessel size (**A**), lumen size (**B**) and W/L ratio (**C**). Panel **D** shows the relation between W/L ratio before dissection and lumen size expansion rate.

Since any single or combined parameters failed to predict the actual lumen size at 45 mm Hg, we sought to construct multivariate binary logistic regression models to check the appropriateness of the artery, in terms of lumen size at the pressurized state. For the prediction of the upper limit (240 μm), the vessel size and the lumen size were selected as the explanatory variables. The model showed 78.8% overall accuracy with 0.25 sensitivity and 0.95 specificity. The odds ratio of the vessel size was 0.913 per unit increase (*P*<0.001) and 1.057 per unit increase of lumen size (*P* = 0.01). The area under the ROC curve was 0.761 (**Figure 6A**). For the prediction of lower limit (160 μm), the vessel size and the W/L ratio before dissection were selected as the explanatory variables. The model showed 92.5% overall accuracy with 0.1 sensitivity and 1.0 specificity. The odds ratio of the vessel size was 1.047 per unit increase (*P*<0.001) and 13.286 per unit increase of the W/L ratio (*P*= 0.042). The area under the ROC curve was 0.747 (**Figure 6B**).

**Figure 6.**
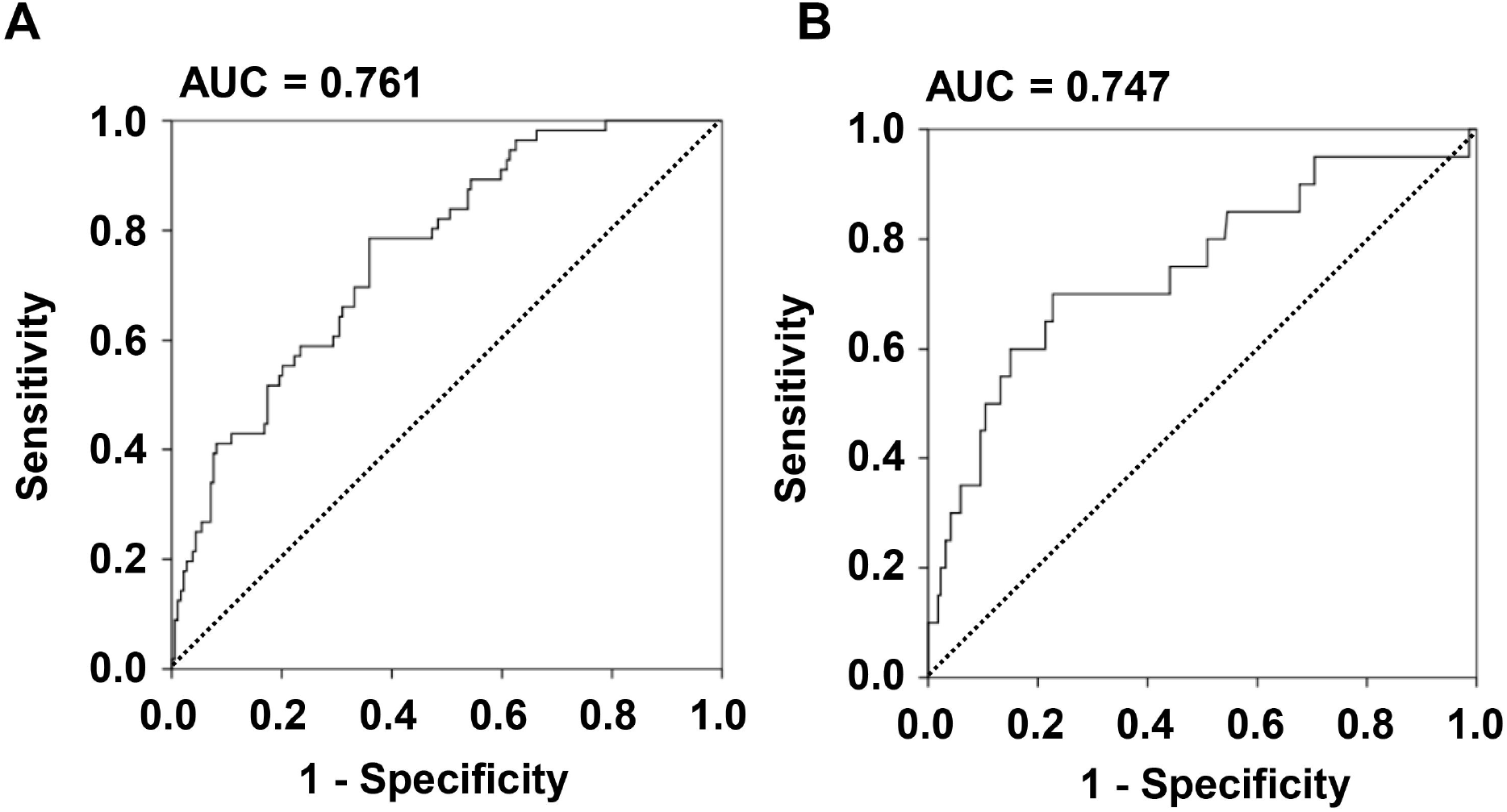
A receiver operator characteristic curves for the binary logistic regression model for vessel selection for myography study. A diagonal dash line represents chance classification. The ROC curves using the threshold of 240 μm (**A**) and 160 μm (**B**) were shown. AUC, area under the curve.

## 4. DISCUSSION

The prognostic value of the structural alteration of small resistance arteries for cardiovascular disease was first reported by Rizzoni et al. in 2003 (3). To date, there have been numerous studies using rodent mesenteric arteries aiming to elucidate the mechanisms of vascular disease. Pressurized myography is one of the most commonly used techniques in vascular research to assess endothelial and smooth muscle function and structural remodeling of small vessels. We have reported for example on the role of immune cells in experimental hypertension using this technique (7). Despite the widespread use and refinement of this method, there remain inefficiencies in the selection of the arteries within the spread mesentery artery tree to study. In the conventional myography protocol, the vessel size becomes clear only after it is mounted on the glass cannulae (8). If the artery size is too small or too large (or if one has actually dissected a vein), the vessel has to be discarded and the same steps repeated from the beginning. This can affect the freshness of the tissue and/or quality of the entire experiment. Although some real-time video tracking systems has been proposed (23), their use is limited to the post-preparation phase and does not allow artery and vein discrimination. Recently, an *in vivo* vessel identification method by two-photon excitation elastin autofluorescence (TPEA) has been proposed (24). These authors showed a clear difference in the signal pattern of the vessel wall between artery and vein, but their method still did not satisfy the need for automatic vessel delineation and size measurement, and requires an expensive two-photon microscope.

U-Net is one of the most successful neural network architectures in the biomedical image segmentation world, used for example in cell counting and morphometry (20,25). Hemelings et al. applied U-Net to a retinal image dataset and achieved high prediction accuracy for artery/vein discrimination, but they did not attempt size measurement (10). A previous study has reported the combination of vessel segmentation and automatic width measurement in retinal images (26). Although this saliency-based approach is efficient for this domain, it does not differentiate between wall and lumen, let alone vein and artery. The potential use of U-Net in vessel size estimation has been recently demonstrated by Stember et al., who used U-Net to predict cerebral aneurysmal area, but the *R*-value between predicted size and radiologist’s manual measurement only reached 0.67 (11).

In this study, we employed U-Net for vessel segmentation followed by automated size measurement with a rectangular approximation algorithm. Our model achieved acceptable performance for vessel segmentation, in terms of IoU and Dice coefficient score. We then applied our size estimation method to the segmented output images. The rectangular-approximation method (using OpenCV) is well validated for manual measurement and also has several advantages over the conventional manual measurement. Most importantly, it is not affected by human perception and therefore free from selection bias. In addition, it provides average diameters in the captured region, which reflect more of the vessel shape, such as bifurcation and branching.

Lastly, we compared vessel size before dissection and after pressurization in a myograph chamber. We found that the vessel sizes are not the same before and after dissection, even at the lowest intraluminal pressure. There are several possible reasons explaining the change in the lumen size after pressurization of the arteries. First of all, the longitudinal strain in the vessel wall will be lost when it is cut; the vessel is no longer axially stretched at the time of analysis. Second, the blood in the lumen will be replaced by physiological salt solution when mounted in the myography chamber, which could affect myogenic tone. Third, the experimental temperature could affect the degree of arterial contraction (27,28). As we collect the mesenteric artery in ice-cold solution to prevent tissue degradation, a certain degree of vasoconstriction could have occurred. It is also possible that our handling with sharp tweezers could induce vasoconstriction, leading to unwanted lumen narrowing and wall thickening.

In contrast to the lumen size itself, its expansion rate has a strong linear correlation with the W/L ratio before dissection. It is interesting but may seem counterintuitive that an artery with higher W/L ratio expands better, because higher W/L ratio usually means vascular remodeling under pressurized conditions (29). However, the relatively thick wall and narrow lumen before dissection is mostly due to constriction and could result in increased capacity to expand in response to positive intraluminal pressure. In other words, the lumen size at certain equilibration pressures can be predictable before mounting in the chamber. Although any single or combined parameters could not directly predict the actual lumen size at 45mmHg, the multivariate binary logistic regression models could facilitate the selection of appropriate vessels for pressurized myography experiments. This finding would be advantageous to investigators, who have been working until now without certainty about the vessel size after mounting.

In summary, this novel approach using U-Net can successfully differentiate 2^nd^ and 3^rd^ order mesenteric arteries from veins and capillaries, and provide vessel segmentation with outer and inner contours, which can then be used for automatic width calculation. The present data-driven approach demonstrates that the lumen diameter at 45 mm Hg of intraluminal pressure can be estimated from the parameters before dissection. We believe that this method streamlines the experimental procedure and may facilitate cardiovascular research.

### LIMITATIONS

There are several limitations to the method we describe. First, the artery/vein discrimination was conducted based on visual and handling assessment, and not confirmed by histological analysis. Therefore, the possibility of mislabeling during the segmentation process cannot be excluded. However, a clear separation of the three layers of the arterial wall in all vessels analyzed with myography system was visible, making this caveat improbable. Second, there is a selection bias with obtained pictures. More “photogenic” arteries were obtained than in fact required in actual experimentation. We tried to minimize this bias by employing 2 investigators to take the mesenteric artery pictures independently (nevertheless based on the same protocol). Third, we could not directly predict the lumen size of arteries pressurized at 45 mm Hg but showed the relation between pre-dissection W/L ratio and fold-change of the lumen size after pressurization. This is partly because of our goal to minimize the number of animals to be euthanized, limiting ourselves to those available from other experiments. An improved prediction model with a larger sample remains to be achieved. We included different experimental mice regardless of age, genotype, and treatment, which could also be a limitation. This is justified by our aim to obtain a universal prediction model for mice with unknown phenotype.

## 5. CONCLUSION

U-Net based segmentation is useful for artery/vein discrimination and automatic size measurement of mouse small mesenteric arteries.

## AUTHOR CONTRIBUTIONS

Akinori Higaki: Conceptualization, Methodology, Software, Formal analysis, Writing-Original draft preparation. Ahmad U. M. Mahmoud: Investigation, Data curation. Pierre Paradis: Validation, Writing-Reviewing and Editing. Ernesto L. Schiffrin: Supervision, Funding acquisition, Writing-Reviewing and Editing.

## Acknowledgement

We are grateful to Adriana Cristina Ene, Isabelle Miguel, and Véronique Michaud for excellent technical support.

