## Supplementary data for "Automated Detection and Diameter Estimation for Mouse Mesenteric Artery using Semantic Segmentation"

**Supplementary Table** Characteristics of mice used for imaging of arteries before dissection and after pressurization at 45 mmHg

|  |  | **Imaging within spread mesenteric artery tree** | | | | | |  | **Imaging at 45 mm Hg** | |
| --- | --- | --- | --- | --- | --- | --- | --- | --- | --- | --- |
| **Strain** | **Genotype** | **# of mice (%)** | **# of images (%)** | **# of Ang II treated-mice (%)** | **TAM (%)** | **Age (days)** | **Male (%)** |  | **# of mice (%)** | **# of images (%)** |
| C57BL/6J | WT | 23 (39) | 244 (37) | 6 (60) | 0 | 83.1 ± 3.9 | 100 |  | 14 (40) | 101 (42) |
| C57BL/6J | *P2rx7^-/-^* | 8 (14) | 80 (12) | 4 (40) | 0 | 91.6 ± 1.8 | 100 |  | 2 (6) | 10 (4) |
| C57BL/6NHsd | ieCre | 7 (12) | 61 (9) | 0 (0) | 57 | 118 ± 19 | 57 |  | 7 (20) | 53 (22) |
| C57BL/6J | *Il-23r^GFP/GFP^* | 5 (8) | 22 (3) | (0) | 0 | 86.0 ± 7.0 | 100 |  | 5 (14) | 21 (9) |
| C57BL/6NHsd | WT | 3 (5) | 33 (5) | 0 (0) | 0 | 143 ± 41 | 67 |  | 2 (6) | 19 (8) |
| C57BL/6NHsd | eET-1 | 2 (3) | 38 (6) | 0 (0) | 0 | 181 ± 0.5 | 100 |  | 0 (0) | 0 (0) |
| C57BL/6NHsd | eET-1/*Apoe^-/-^*/*Nox4*^+/-^ | 2 (3) | 24 (4) | 0 (0) | 0 | 151± 2.0 | 100 |  | 2 (6) | 21 (9) |
| C57BL/6NHsd | ieET-1 | 2 (3) | 5 (1) | 0 (0) | 100 | 169 ± 0 | 100 |  | 2 (6) | 5 (2) |
| C57BL/6NHsd | eET-1/*Apoe^-/-^* | 1 (2) | 35 (5) | 0 (0) | 0 | 92 | 100 |  | 0 (0) | 0 (0) |
| C57BL/6NHsd | *Apoe^-/-^* | 1 (2) | 27 (4) | 0 (0) | 0 | 97 | 100 |  | 0 (0) | 0 (0) |
| C57BL/6NHsd | eET-1/*Apoe*^+/-^/*Nox1*^-/y^ | 1 (2) | 23 (4) | 0 (0) | 0 | 116 | 100 |  | 0 (0) | 0 (0) |
| C57BL/6NHsd | *Apoe*^+/-^/*Nox1*^-/y^ | 1 (2) | 20 (3) | 0 (0) | 0 | 115 | 100 |  | 0 (0) | 0 (0) |
| C57BL/6NHsd | *Apoe^-/-^*/*Nox1*^-/y^ | 1 (2) | 16 (2) | 0 (0) | 0 | 128 | 100 |  | 0 (0) | 0 (0) |
| C57BL/6NHsd | eET-1/*Apoe^-/-^*/*Nox1*^-/y^ | 1 (2) | 16 (2) | 0 (0) | 0 | 128 | 100 |  | 0 (0) | 0 (0) |
| C57BL/6NHsd | *ApoE^-/-^*/*Nox4*^+/-^ | 1 (2) | 10 (2) | 0 (0) | 0 | 147 | 100 |  | 1 (3) | 10 (4) |

Data are shown as number (#) and % of mice and images and as means ± SEM for ages. Ang II, angiotensin II; *Apoe^-/-^*, apolipoprotein E knockout; ieCre, tamoxifen-inducible endothelium-restricted Cre recombinase transgenic; ieET-1, tamoxifen-inducible endothelium-restricted human endothelin-1 overexpressing transgenic; eET-1, endothelium-restricted human endothelin-1 overexpressing transgenic; *Il-23r^GFP/GFP^*, interleukin-23 receptor green fluorescent protein knock-in; *Nox1^+/-^* and *Nox1^-/y^*, NADPH oxidase 1 heterozygote and knockout; *Nox4^-/-^*, NADPH oxidase 4 knockout; *P2rx7^-/-^*, purinergic receptor P2X purinoceptor7 knockout; WT, wild-type.

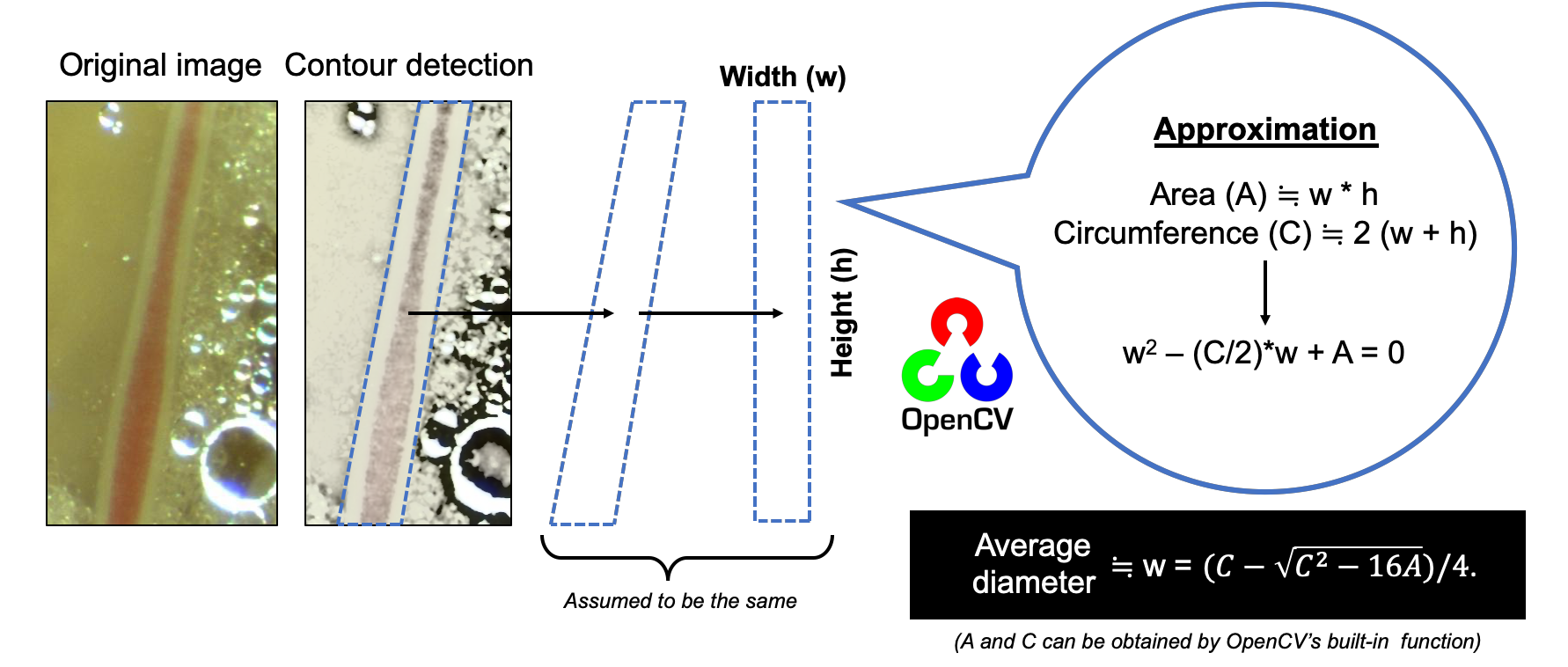

**Supplementary Figure. S1. Rectangular approximation method to estimate the average diameter of the extracted contour.** Vessel contours are extracted by the U-net segmentation model. The contour has an elongated, wavy trapezoidal shape that can be approximated to a rectangular that has the same area and circumference.

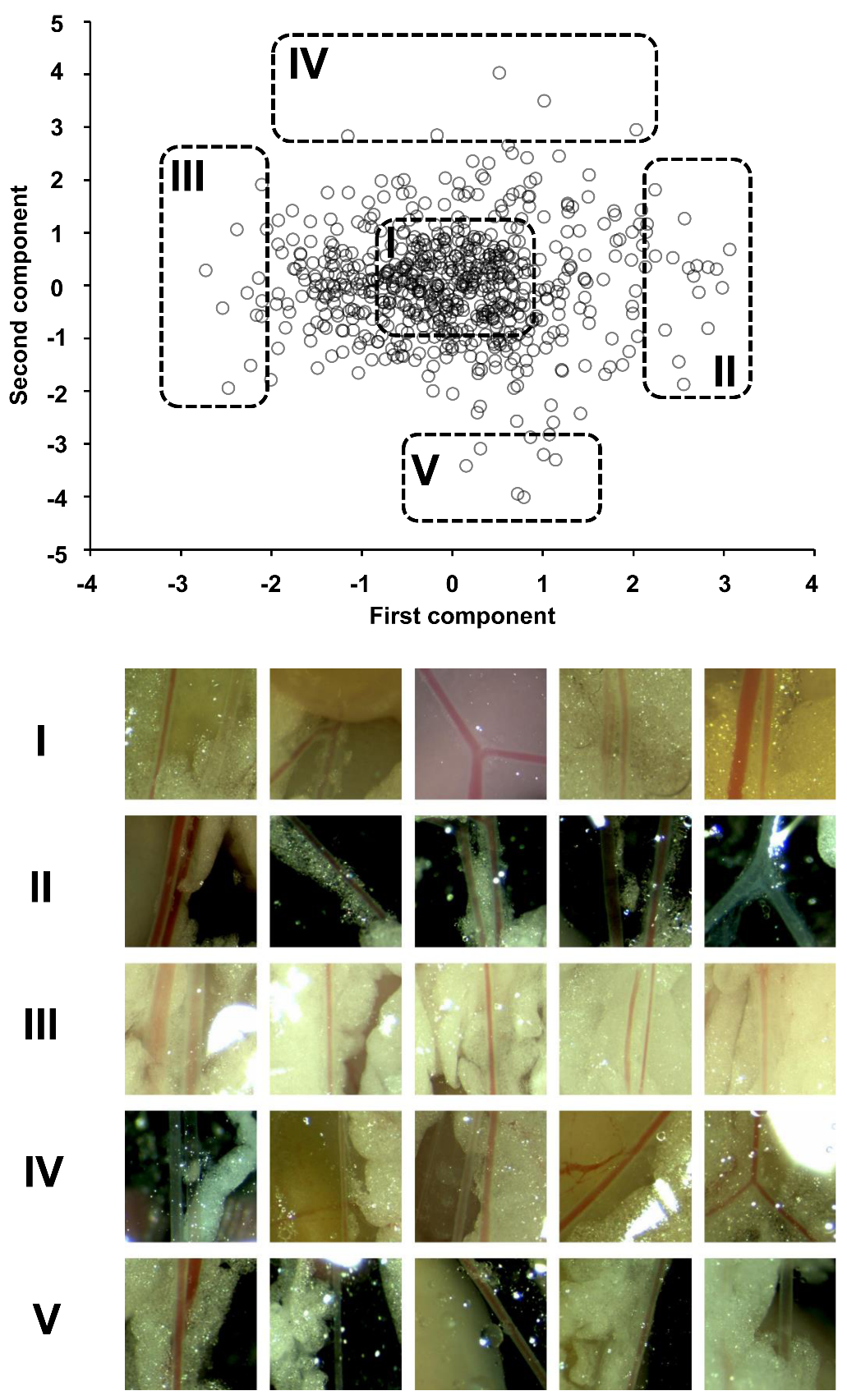
**Supplementary Figure. S2** **Examples of collected images visualized with the principal components.** Each data point in the scatterplot represents a microscopic in situ image within the spread mesenteric artery tree used for the modeling. According to the principal component analysis, five representative images in the specific plot area (from I to V) are shown in the bottom panel.

**Supplementary Figure. S3** Correlation between manual measurement and the automated calculation with rectangular approximation. The linear correlation between manual measurement and automatic calculation for vessel diameter (A) and lumen diameter (B) is shown.

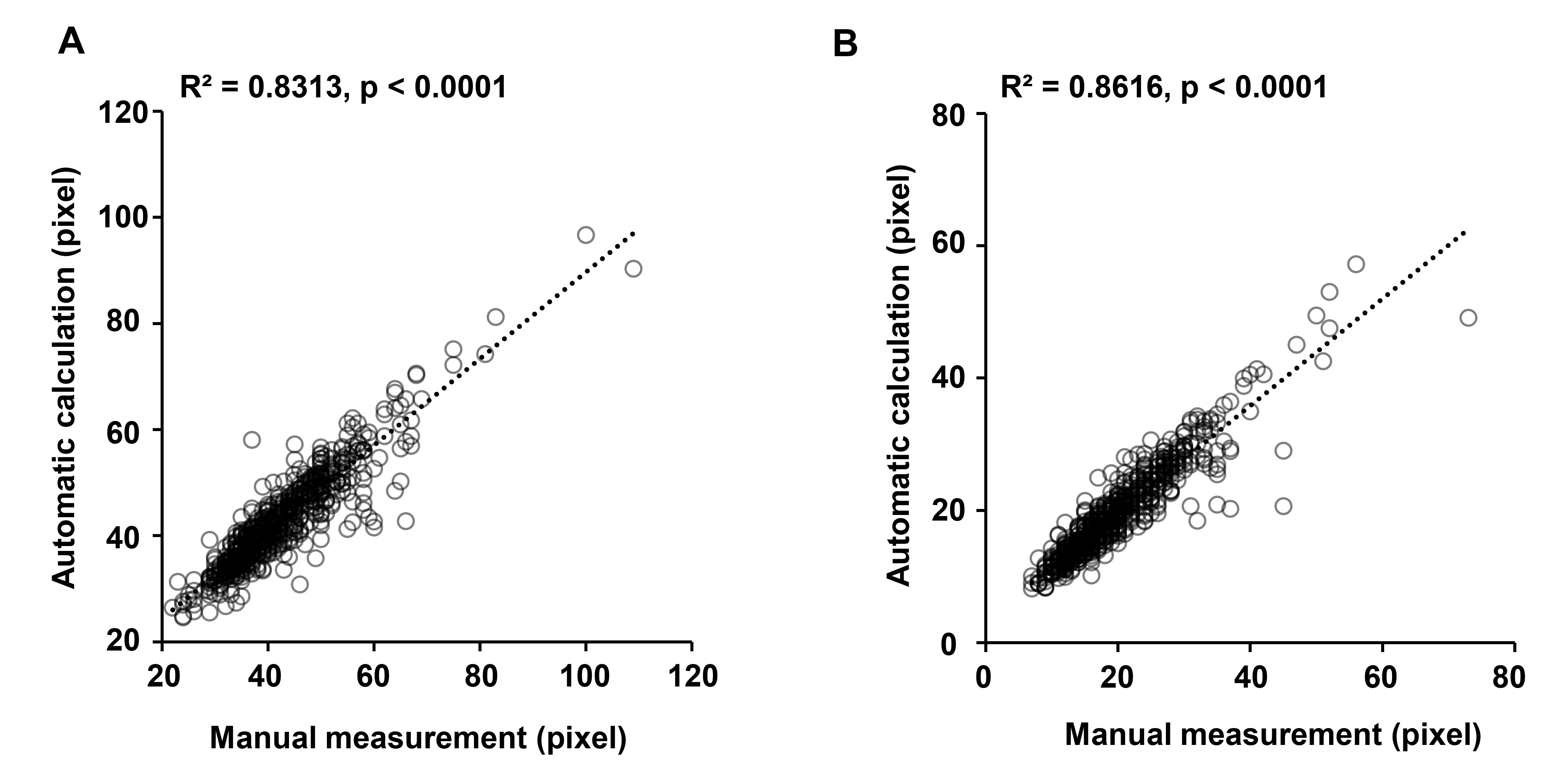

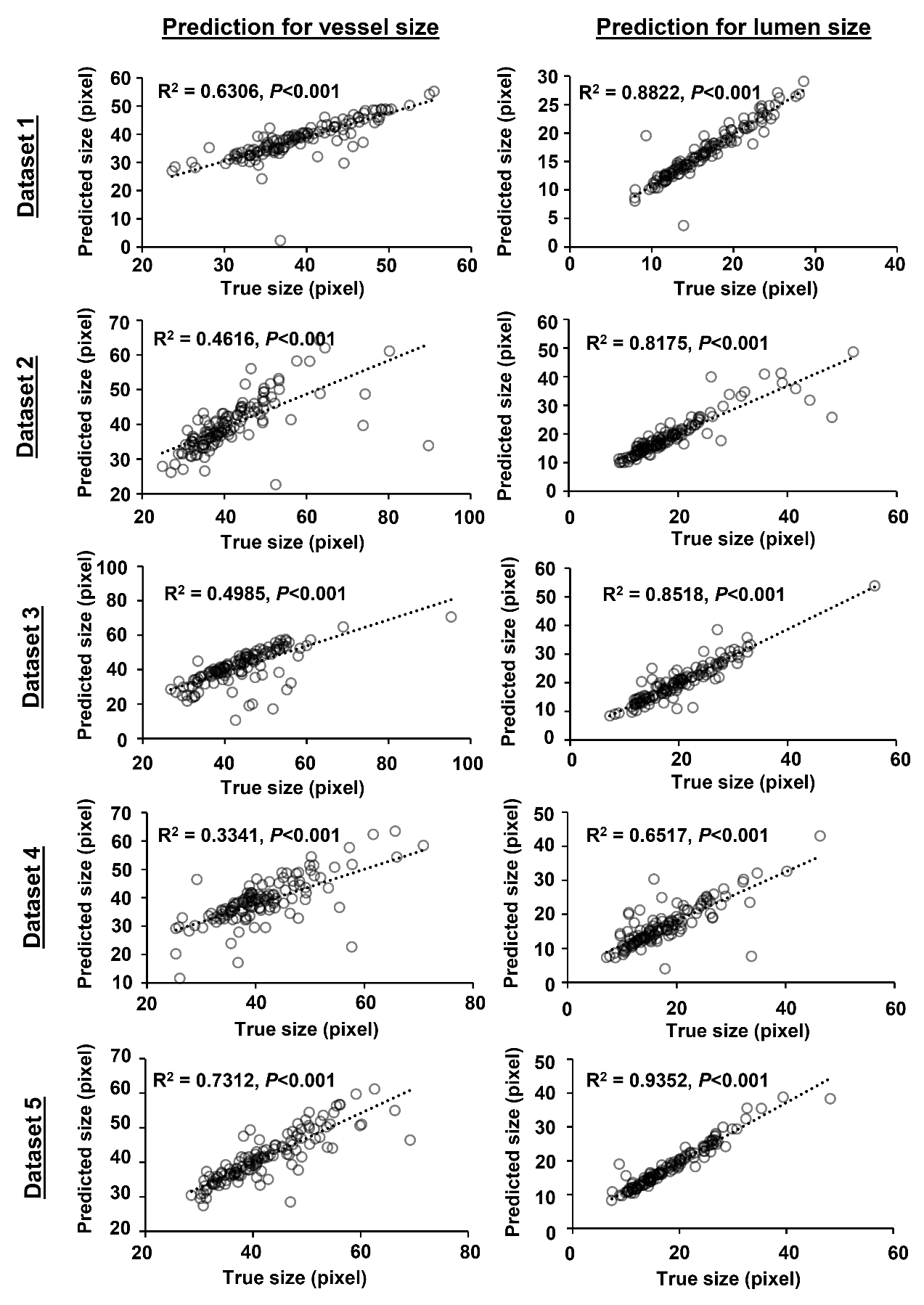
**Supplementary Figure. S4** Correlation of the vessel sizes between true contour-based and the predicted contour-based calculation. According to the datasets from the 5-fold cross validation, 5 scatter plots were shown for the vessel size (left column) and for the lumen size (right column) respectively.
